# Impact of circadian time of dosing on cardiomyocyte-autonomous effects of glucocorticoids

**DOI:** 10.1101/2021.12.30.474468

**Authors:** Michelle Wintzinger, Manoj Panta, Karen Miz, Ashok D. Pragasam, Hima Durumutla, Michelle Sargent, Clara Bien Peek, Joseph Bass, Jeffery D. Molkentin, Mattia Quattrocelli

**Author notes:** These Authors contributed equally (Michelle Wintzinger, Manoj Panta). **Corresponding author** - Mattia Quattrocelli, PhD, Molecular Cardiovascular Biology, Heart Institute, Cincinnati Children’s Hospital Medical Center, Dept. Pediatrics – UCCOM, 240 Albert Sabin Way T4.676, Cincinnati, OH 45229. **Data availability** - RNA-seq datasets reported here are available on GEO with accession number GSE186875. All data and analyses needed to evaluate the conclusions in the paper are present in the paper and/or the Supplementary Materials.

## Abstract

Mitochondrial capacity is critical to adapt the high energy demand of the heart to circadian oscillations and diseased states. Glucocorticoids regulate the circadian cycle of energy metabolism, but little is known about how circadian timing of exogenous glucocorticoid dosing directly regulates heart metabolism through cardiomyocyte-autonomous mechanisms. While chronic once-daily intake of glucocorticoids promotes metabolic stress and heart failure, we recently discovered that intermittent once-weekly dosing of exogenous glucocorticoids promoted muscle metabolism in normal and obese skeletal muscle. However, the effects of glucocorticoid intermittence on heart metabolism and heart failure remain unknown. Here we investigated the extent to which circadian time of dosing regulates the effects of the glucocorticoid prednisone in heart metabolism and function in conditions of single pulse or chronic intermittent dosing. In WT mice, we found that prednisone improved cardiac content of NAD^+^ and ATP with light-phase dosing (ZT0), while the effects were blocked by dark-phase dosing (ZT12). The drug effects on mitochondrial function were cardiomyocyte-autonomous, as shown by inducible cardiomyocyte-restricted glucocorticoid receptor (GR) ablation, and depended on an intact cardiomyocyte clock, as shown by inducible cardiomyocyte-restricted ablation of Brain and Muscle ARNT-like 1 (BMAL1). Conjugating time-of-dosing with chronic intermittence, we found that once-weekly prednisone improved metabolism and function in heart after myocardial injury dependent on circadian time of intake, i.e. with light-phase but not dark-phase dosing. Our study identifies cardiac-autonomous mechanisms through which circadian-specific intermittent dosing reconverts glucocorticoid drugs to metabolic boosters for the heart.

## INTRODUCTION

Bioenergetic capacity is critical to adapt the high energy demand of the heart to circadian oscillations and/or diseased states. Endogenous glucocorticoids are conserved pleiotropic hormones regulating the circadian cycle of energy utilization and storage in virtually all our organs [1]. However, little is known about the extent to which circadian timing of exogenous glucocorticoid dosing impacts function of the primary receptor of these drugs, the glucocorticoid receptor (GR). The question of time-restricted pharmacology is critical for the heart, which displays circadian oscillations in nutrient metabolism [2]. The circadian clock is important for heart function, as shown by the fact that genetic ablation of BMAL1, core component of the activating clock complex in every tissue, disrupts heart metabolism and function [3]. However, mechanisms and relevance of interplay between circadian time of glucocorticoid dosing and heart metabolism remain unaddressed.

The GR regulates and is in turn regulated by core components of the circadian clock [1; 4]. The autonomous circadian clock relies on the permutation between the activating complex, e.g. CLOCK/BMAL factors active during the light-phase, and the repressing complex, e.g. PER/CRY factors active during the dark-phase [5]. In mice, endogenous glucocorticoids entrain the circadian rhythm of GR activation by peaking during the dark phase and dipping during the light-phase [6]. Indeed, GR is known to regulate nocturnal factors like PER2 [1]. In turn, the clock factors Cry1/2 and CLOCK inhibit GR activity during the dark-phase [7] and during the late light-phase [8], respectively. This leads to a critical time window in the early light-phase (corticosterone trough [6]), where the GR is not repressed and not occupied by endogenous steroids, thus highly responsive to exogenous glucocorticoids [7; 9]. However, these molecular regulations were found in noncardiac cells and therefore the effects of light-phase-restricted synthetic glucocorticoids (anti-physiologic to endogenous glucocorticoids) remain unknown for heart function and metabolism.

The glucocorticoid receptor (GR) is important for normal heart function and contractility [10; 11]. Indeed, GR activation by exogenous glucocorticoids improves calcium handling in cardiomyocytes [10; 12]. However, the cardiac-autonomous effects of GR pharmacology on heart metabolism are still unknown. This question is particularly important as the heart-autonomous versus body-wide effects of chronic glucocorticoid intake on cardiac health remain unresolved. Synthetic glucocorticoids are used chronically by >2.5M people in the US [13], but their regular once-daily intake promotes metabolic syndrome and cardiovascular diseases in a dose-dependent manner [14]. Opposite to once-daily dosing, we discovered that intermittent dosing (once-weekly) of glucocorticoids improves function and mitochondrial capacity in skeletal muscle in normal and metabolic stress conditions[15; 16]. Dosing intermittence could therefore reverse the adverse benefits/risks ratio of chronic steroid regimens for the heart. However, the cardiac effects and cardiomyocyte-autonomous mechanisms of glucocorticoid intermittence are still unknown.

Here we investigated the extent to which circadian time of dosing regulates the cardiac effects of the exogenous glucocorticoid prednisone in conditions of single pulse or chronic intermittent dosing. In WT mice, we found that prednisone improved NAD^+^ and ATP levels when dosed at the start of the light-phase, while these effects were blocked with dosing at the start of the dark-phase. The effects on mitochondrial function depended on cardiomyocyte-specific GR, as shown by inducible cardiomyocyte-specific GR ablation, and on an intact cardiomyocyte-autonomous clock, as shown by inducible cardiomyocyte-specific BMAL1 ablation. Conjugating time-of-dosing with chronic intermittence, we found that once-weekly light-phase prednisone stimulated a transcriptional program relevant for circadian metabolism and promoted function in the failing heart after myocardial infarction. Our study identifies cardiac-autonomous mechanisms through which circadian time and chronic intermittence reconvert glucocorticoid drugs to bioenergetic boosters for the heart.

## RESULTS

### Pro-metabolic effects of prednisone in myocardium depend on time-of-dosing

We sought to investigate the acute effects of exogenous glucocorticoids on heart metabolism according to circadian time of dosing. To this end, we treated WT mice (C57BL/6 background; 8 weeks of age) with a single injection of i.p. 1 mg/kg prednisone at either ZT0 (start of light-phase) or ZT12 (start of dark-phase). Drug dose and times of dosing were consistent with our previous chrono-pharmacology study in skeletal muscle [17]. We assayed metabolites and gene expression in myocardial tissue every 4 hours starting at 1-hour post-pulse and during one circadian cycle post-pulse. For metabolite quantitation, we focused our analyses on myocardial NAD^+^ and ATP content, quantitating these metabolites through targeted assays as metabolic biomarkers of mitochondrial function. NAD^+^ is a fundamental determinant and biomarker of mitochondrial health in a variety of cardiovascular conditions [18]. ATP is produced by oxidative phosphorylation in mitochondria and the ATP synthase is directly regulated by the NAD^+^ redox balance in heart [19]. As parallel analyses of related gene expression, we monitored the effects of time-specific prednisone on myocardial expression of *Nampt* and *Ppargc1a. Nampt* encodes the homonymous rate-limiting enzyme for NAD^+^ biogenesis [20]. *Ppargc1a* encodes the mitochondrial regulator PGC1α, which promotes ATP production in heart through mitochondrial biogenesis and function [21].

Compared to vehicle, the light-phase prednisone pulse increased the levels of NAD^+^ and ATP content **(Fig. 1A)**, as well as *Nampt* and *Ppargc1a* expression levels **(Fig. 1B)** over the course of the post-pulse circadian cycle. However, these effects were blunted or absent after dark-phase prednisone pulse **(Fig. 1A-B)**. We monitored clock gene expression to check for possible regimen-specific changes in heart. We found that the light-phase prednisone pulse – but not the dark-phase pulse – increased amplitude of activating clock complex genes *Arntl* (gene name for BMAL1) and *Clock* without major changes in their rhythmicity. Conversely, we found that the dark-phase pulse – but not the light-phase pulse – transiently increased repressive clock complex genes *Per1* and *Cry2* during their normal trough phase **(Fig. 1C)**.

**Figure 1.**
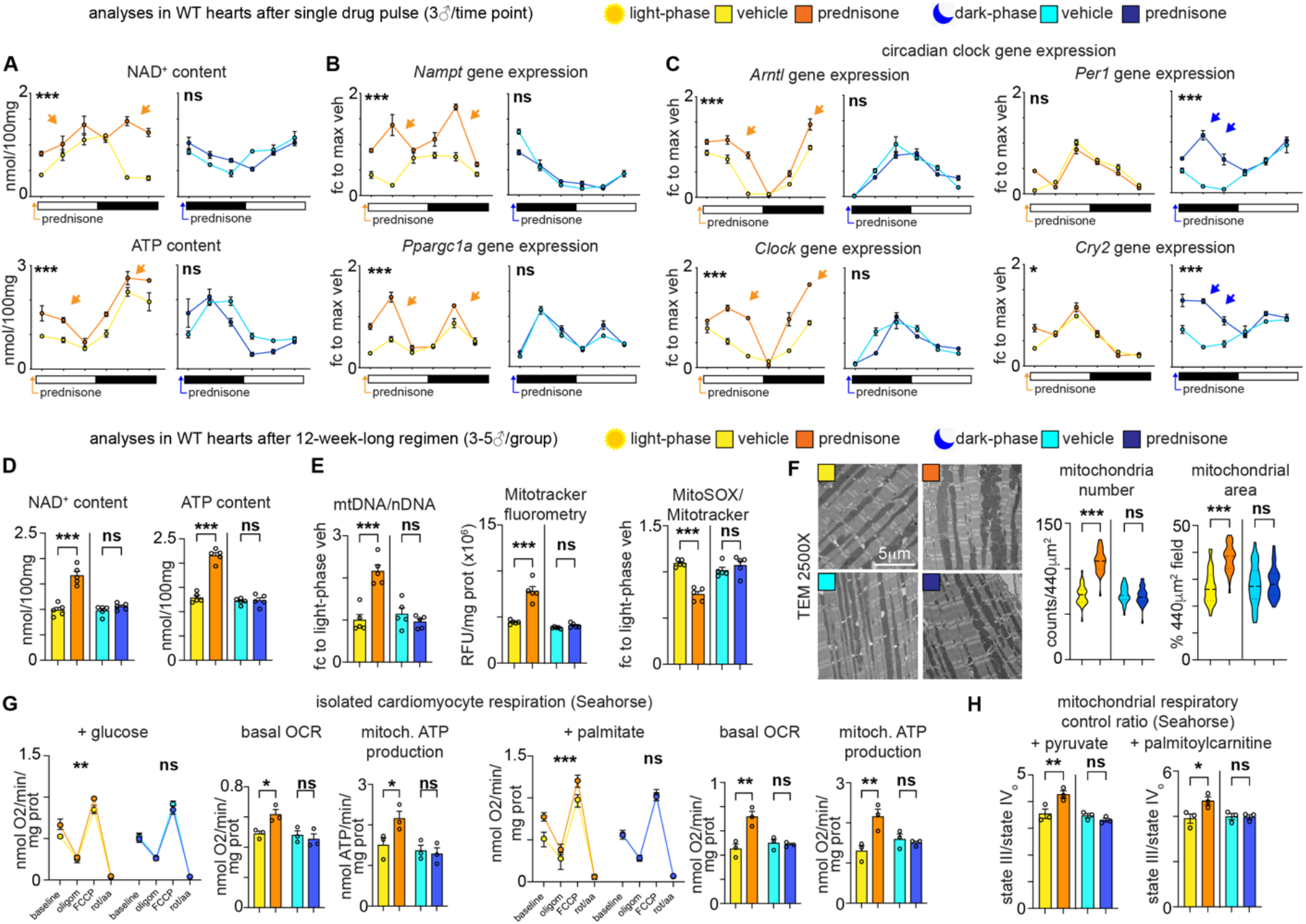
Light-phase but not dark-phase injection of pulsatile or intermittent prednisone increases biomarkers of mitochondrial metabolism in the myocardium. **(A-B)** Mice were injected with one dose of 1mg/kg i.p. prednisone in vivo and sampled every 4 hours from 1-hour post-injection. Light-phase prednisone (pulse at ZT0) increased NAD^+^ and ATP content **(A)**, as well as *Nampt* and *Ppargc1a* expression **(B)**, in hearts in vivo in two circadian phases following injection (arrows). These effects were blocked with a dark-phase drug pulse (ZT12). **(C)** The lightphase prednisone pulse correlated with increased amplitude of *Arntl* (BMAL1) and *Clock* expression (activating clock complex), while the dark-phase prednisone transiently elevated *Per1* and *Cry2* gene expression (repressive clock complex) during their circadian trough. **(D-H)** Mice were injected with once-weekly 1mg/kg prednisone for 12 weeks at either ZT0 (light-phase treatment) or ZT12 (dark-phase). Compared to isochronic vehicle treatments, light-phase but not dark-phase treatments improved NAD^+^ and ATP levels in myocardium **(D)**. This correlated with increased mitochondrial abundance, estimated by mtDNA/nDNA ratio and Mitotracker fluorometry in tissue, and decreased ROS levels, estimated through MitoSOX fluorometry **(E)**. TEM imaging showed increased mitochondria number and mitochondria-filled areas in light-phase-but not dark-phase-treated myocardial sections **(F)**. Seahorse respirometry in isolated cardiomyocytes showed increased glucose-or lipid-fueled basal respiration and ATP production after light-phase but not dark-phase treatment **(G)**. These trends correlated with RCR trends in isolated mitochondria with either pyruvate or palmitoylcarnitine **(H)**. * = p < 0.05, ** = p < 0.01, *** = p < 0.001; 2w ANOVA (curves in A-B, G), 1w ANOVA + Sidak (D-H).

We then asked whether the metabolic changes of light-phase prednisone in heart were maintained with a chronic treatment. Regarding chronic steroid dosing, we used the 12-week-long once-weekly 1mg/kg prednisone intermittent dosing that we successfully tested with circadian-specific effects in WT skeletal muscle [17]. Intermittent prednisone dosing promotes a pro-metabolic signature, reversing the obesogenic effects induced by once-daily dosing [16]. We treated WT mice with 12-week-long intermittent prednisone regimens through dosing at either ZT0 (light-phase treatment) or ZT12 (dark-phase treatment). At treatment endpoint and compared to isochronic (time-matched) vehicle controls, light-phase treatment increased NAD^+^ and ATP content in the myocardium, while these effects were blocked by dark-phase treatment **(Fig. 1D)**. Considering the upregulation of PGC1α expression, we quantitated mitochondrial abundance in the myocardium through parallel analyses with mitochondrial DNA ratio over nuclear DNA (mtDNA/nDNA) [22]; unbiased fluorometry in myocardial tissue for Mitotracker Green FM (mitochondrial signal independent from mitochondrial membrane potential) and MitoSOX Red (reactive oxygen species (ROS) signal) [23]; transmission electron microscopy (TEM) imaging [24; 25]; protein analysis of mitochondrial marker TOMM20 [26]. Light-phase treatment increased mtDNA/nDNA ratio and normalized Mitotracker signal in myocardium, while decreasing MitoSOX signal **(Fig. 1E)**. These data were in accordance with gains in mitochondrial number and mitochondria-occupied areas in TEM images **(Fig. 1F)**, as well as with tissue levels of TOMM20 **(Suppl. Fig. 1A)**. The changes induced by light-phase treatment did not appear dependent on protein level changes in glucocorticoid receptor (GR), or its co-factors NCOR1 (co-repressor) and CREBBP (co-activator) [27; 28], or agonists of mitochondrial fusion/fission MFN2 and PARKIN [29] **(Suppl. Fig. 1B)**.

Furthermore, we measured mitochondrial respiratory function through Seahorse-based respirometry in isolated cardiomyocytes [30], quantitating basal respiration and ATP production in glucose- or palmitate-fueled conditions to monitor for possible shifts in substrate use. To validate that the trends in cardiomyocyte respiration were due to intrinsic changes in mitochondria, we also performed parallel Seahorse analyses of respiratory control ratio (RCR) in isolated mitochondria. The RCR estimates the capacity of isolated mitochondria to oxidize substrate to phosphorylate ADP and is indeed calculated as ratio of state III (ADP) over state IVo (oligomycin) OCR values [31]. Light-phase treatment increased basal cardiomyocyte respiration and ATP production rate independent from macronutrient fuel **(Fig. 1G)**, correlating with increased mitochondrial RCR levels with either pyruvate or palmitoylcarnitine as substrates **(Fig. 1H)**. Importantly, the treatment effects on mitochondrial metabolites, abundance and respiratory function were blocked with dark-phase dosing **(Fig. 1D-H)**.

Thus, circadian time of intake regulated the acute and chronic effects of the GC prednisone on mitochondrial metabolism and abundance in the normal heart.

### Light-phase intermittent prednisone activates a transcriptional program involving circadian-metabolic pathways in heart and improves function and metabolism after myocardial infarction

We asked whether the effects of light-phase prednisone were significant in the context of myocardial infarction (MI), where mitochondrial capacity is a key target to improve overall cardiac function [32]. We treated sham- or MI-operated WT mice from 2-weeks post-surgery to avoid early changes in pathophysiologic remodeling and provide a comparable control/dysfunction baseline before onset of treatments. MI or sham operations were conducted at 8 weeks of age after baseline echocardiography, and treatments (once-weekly i.p. 1mg/kg prednisone or vehicle at ZT0) started after confirming decreases in fractional shortening and stroke volume at 2-weeks post-MI. Treatment was administered for 6 weeks and analyses were conducted in both sham- and MI-operated mice.

Compared to vehicle at endpoint, light-phase intermittent prednisone treatment significantly increased fractional shortening and stroke volume after MI to levels close to control sham hearts. The trends were not significant in sham-operated hearts **(Fig. 2A)**.

**Figure 2.**
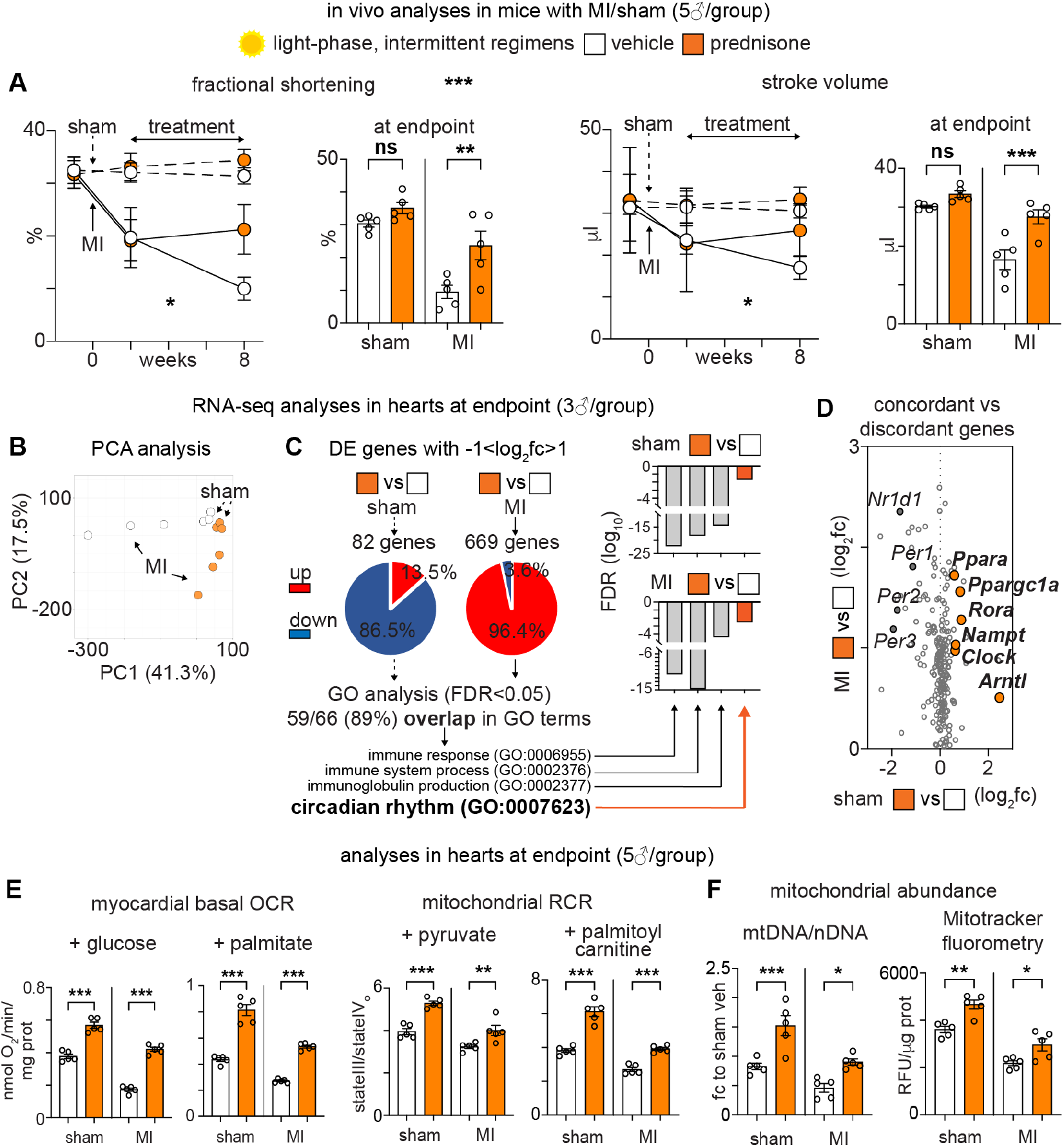
Chronic treatment with light-phase intermittent prednisone improves heart function and metabolism after myocardial infarction. WT mice were randomized into sham or myocardial infarction (MI) cohorts and treated with light-phase intermittent 1mg/kg prednisone for 6 weeks from 2-weeks post-MI. **(A)** Treatment improved fractional shortening and stroke volume in mice with MI. **(B)** PCA analysis of RNA-seq profiles showed treatment-associated clustering in both sham and MI hearts. **(C)** Differentially expressed (DE) genes in treated versus control hearts showed 89% overlap in GO analysis between sham and MI conditions. Among overlapping GO terms were inflammation and circadian rhythm pathways. **(D)** Chart presenting genes with concordant versus discordant regulation by treatment in sham versus MI conditions. Concordant genes are on the right side, while discordant genes are on the left side. Activating clock complex genes *(Arntl, Clock, Rora),* as well as metabolic regulators *Nampt, Ppargc1a* and *Ppara* were among concordant genes. Repressive clock complex genes *(Nr1d1, Per1-3)* were among discordant genes. **(E-F)** Treatment increased mitochondrial respiration (tissue OCR and mitochondrial RCR) and mitochondrial abundance (Mitotracker fluorometry in myocardial tissue and mtDNA quantitation) in both sham and MI hearts. * = p < 0.05, ** = p < 0.01, *** = p < 0.001; 2w ANOVA (curves in A), 1w ANOVA + Sidak (A, E-F); Benjamini-Hochberg for GO term FDR (C).

At endpoint, we analyzed sham-vehicle, sham-treated, MI-vehicle and MI-treated hearts through RNA-seq. Principal component analysis of whole transcriptomes clustered samples according to surgery and treatment **(Fig. 2B)**. We then performed pathway analysis in the genes that were differentially expressed (DE) by treatment in each setting using the arbitrary thresholds of >2-fold change and >10CPM abundance (FDR<0.05). Intriguingly, we found that treatment increased the number of DE genes from 82 in sham hearts to 669 in MI hearts **(Suppl. Table 1)**. Also, the overall trend in treatment-driven regulation in DE genes shifted from 87% downregulated in sham to 96% upregulated in MI hearts **(Fig. 2C)**. Strikingly though, the gene ontology (GO) analysis revealed a high degree of overlap in pathways (89% overlap in GO terms) between DE genes in sham versus MI settings **(Suppl. Table 2)**. Among these overlapping GO terms, we noticed several inflammation-related pathways, as well as circadian rhythm pathways **(Fig. 2C)**. We therefore analyzed concordant and discordant genes for genes contained in the circadian rhythm GO pathways. We defined “concordant genes” as upregulated by treatment in both sham and MI hearts, while “discordant genes” were upregulated by treatment in MI hearts but downregulated in sham hearts (no arbitrary thresholds). Intriguingly, we found that activating clock complex genes *(Arntl, Clock, Rora),* as well as metabolic regulators *Nampt, Ppargc1a* and *Ppara* (cofactor of PGC1α) were among concordant genes, i.e. upregulated by treatment in both sham and MI hearts **(Fig. 2D; Suppl. Fig. 2)**. Conversely, repressive clock complex genes *(Nr1d1, Per1-3)* were among discordant genes, i.e. upregulated by treatment in MI but not in sham hearts **(Fig. 2D)**.

We quantitated mitochondrial respiration in the left ventricle myocardium of sham and MI hearts at endpoint, performing parallel respirometry analyses of basal oxygen consumption rate (OCR) in freshly collected myocardial slices [33] and of RCR in isolated mitochondria. Treatment increased tissue OCR and mitochondrial RCR in sham and MI hearts, rescuing the levels in MI hearts to levels like in control sham hearts **(Fig. 2E)**. We then quantitated changes in mitochondrial abundance, and both mtDNA/nDNA and MitoTracker assays showed gain of mitochondrial density with treatment in sham and MI hearts **(Fig. 2F)**.

Thus, intermittent dosing of light-phase prednisone improved systolic function after MI, correlating with a circadian-metabolic transcriptional program and increased levels of mitochondrial function and abundance in the myocardium.

### Acute transcriptional and metabolic effects of light-phase prednisone in heart require cardiomyocyte-autonomous GR and BMAL1

Considering the high level of concordance between transcriptional effects and metabolic remodeling produced by light-phase prednisone in heart, we investigated the extent to which the cardiomyocyte-autonomous glucocorticoid receptor (GR, encoded by the *Nr3c1* gene) mediates these effects. To this end, we generated inducible heart-specific GR-KO mice crossing *Myh6-CreERT^+/-^* [34] with *Nr3c1^fl/fl^* [35] transgenic lines on the C57BL/6 background. At 8 weeks of age, we induced GR ablation in heart through a sequence of i.p. tamoxifen injections (20mg/kg/day for 5 days) followed by 14 days on tamoxifen-containing chow (40mg/kg) and 2 days of regular chow for tamoxifen washout. Such induction paradigm ablated ~85% of GR in heart without immediate changes in plasma corticosterone or myocardial tissue OCR at baseline **(Suppl. Fig. 3A)**. We compared GR-WT *(Nr3c1^fl/fl^; Myh6-CreERT^-/-^)* versus GR-KO *(Nr3c1^fl/fl^; Myh6-CreERT^+/-^)* littermates for the effects of a light-phase prednisone pulse in vivo immediately after tamoxifen exposure and washout, i.e. at ~12 weeks of age. We injected GR-WT and GR-KO mice with a single i.p. 1mg/kg prednisone dose at ZT0 (light-phase) and monitored the effects in heart at 24-hours postdose. For this experiment, we focused on the single pulse setting and not the chronic regimen setting to avoid possible confounding effects of long-term GR ablation in heart [36].

Following our analyses in WT myocardium, we quantitated mitochondrial respiration, NAD^+^ and ATP levels, and *Nampt* and *Pparcg1a* expression. In GR-WT mice the drug pulse recapitulated the gains in both basal myocardial OCR and mitochondrial RCR as compared to vehicle regardless of substrate, while these effects were blocked in GR-KO hearts **(Fig. 3A)**. This correlated with analogous trends in NAD^+^ and ATP content in the left ventricle myocardial tissue **(Fig. 3B)**. Intriguingly, inducible GR ablation blocked the transcriptional effects of the drug pulse on *Nampt* and *Ppargc1a* in heart **(Fig. 3C)**.

**Figure 3.**
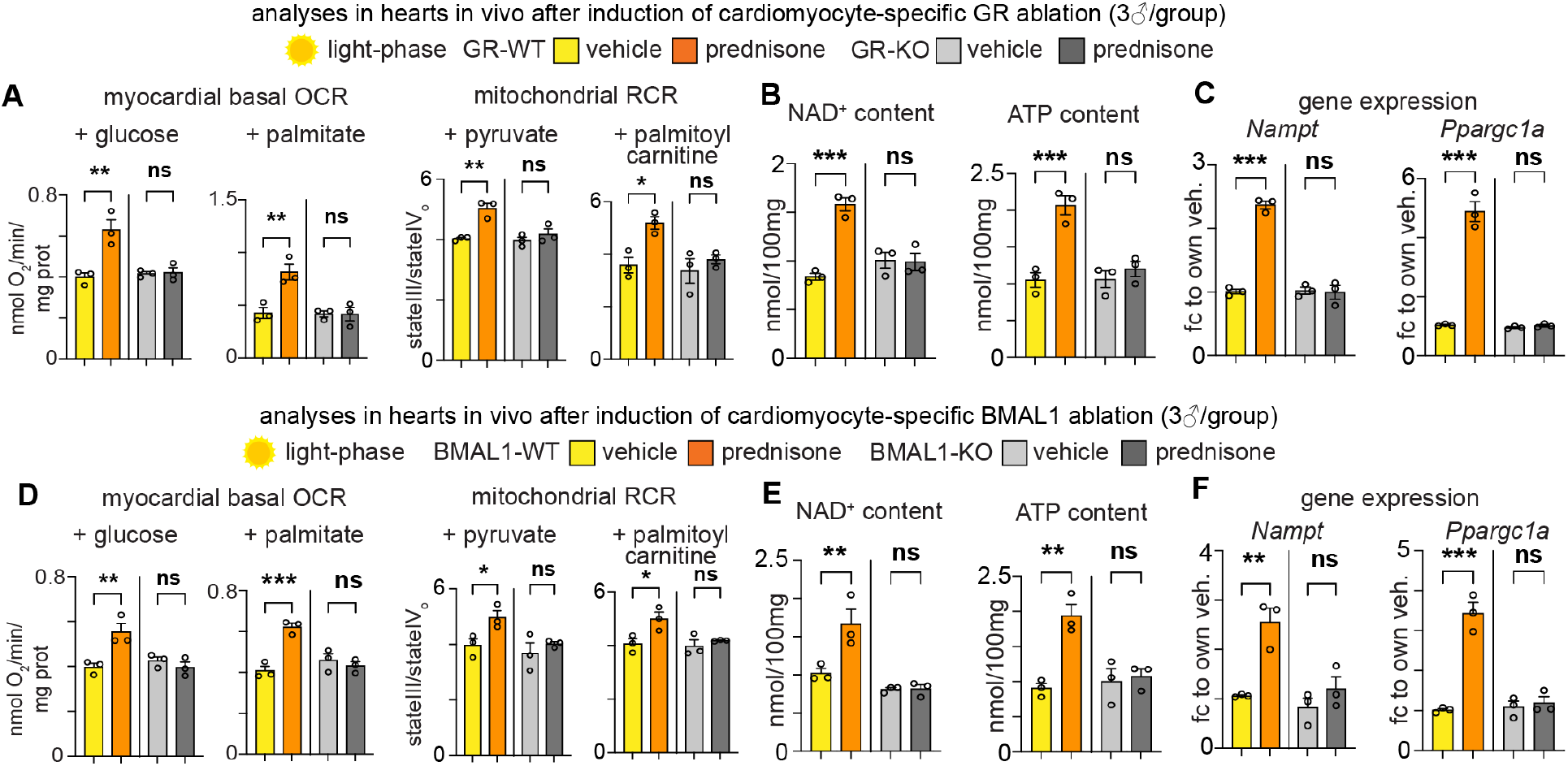
The mitochondrial effects of light-phase prednisone are dependent upon cardiomyocyte-specific GR and BMAL1 expression. **(A)** At 24-hours post-injection, light-phase prednisone increased mitochondrial respiration in GR-WT but not GR-KO hearts (induced cardiomyocyte-specific GR ablation in postnatal tissue), as compared to respective vehicle controls. **(B-C)** Analogous trends were found in myocardial content of NAD^+^ and ATP, as well as expression levels of *Nampt* and *Ppargc1a.* **(D-F)** Analogous trends in myocardial basal respiration, RCR in isolated mitochondria, NAD^+^ and ATP content, and *Nampt* and *Ppargc1a* expression were found comparing BMAL1-WT to BMAL1-KO hearts (induced cardiomyocyte-specific BMAL1 ablation in postnatal tissue). * = p < 0.05, ** = p < 0.01, *** = p < 0.001; 1w ANOVA + Sidak.

Thus, the cardiomyocyte-autonomous GR is required for the transcriptional and metabolic effects of light-phase prednisone in heart.

Furthermore, we were intrigued by the concordant trends induced by light-phase intermittent prednisone on cardiac expression of genes encoding factors of the activating clock complex. We thus asked whether an intact cardiomyocyte-autonomous activating clock complex was required for the cardiac effects of light-phase prednisone. Indeed, the activating clock complex, which encompasses BMAL1, is normally active during the light-phase in mice [5]. Importantly, BMAL1 (gene name, *Arntl)* promotes mitochondrial function and NAD^+^ biogenesis [37]; plays a key role in cardiomyocyte metabolism and function [3]; regulates the epigenetic activity of GR in another striated muscle context, the skeletal muscle [17]. We therefore asked whether cardiomyocyte-specific deletion of BMAL1 abrogated the effects of light-phase prednisone in heart. To this end, we generated inducible heart-specific BMAL1-KO mice crossing *Myh6-CreERT^+/-^* [34] with *Arntl^fl/fl^* [38] transgenic lines on the C57BL/6 background. We promoted gene ablation following age and tamoxifen settings as aforementioned. We attained >90% ablation in heart with no immediate baseline changes in plasma corticosterone or myocardial OCR **(Suppl. Fig. 3B)**. We compared BMAL1-WT *(Arntl^fl/fl^; Myh6-CreERT^-/-^*) versus BMAL1-KO (*Arntl^fl/fl^; Myh6-CreERT^+/-^*) littermates for the effects of a light-phase prednisone pulse in vivo (ZT0) immediately after tamoxifen exposure and washout, i.e. at ~12 weeks of age. Also for this experiment, we focused on the single pulse setting and not the chronic regimen setting to avoid significant changes in overall heart function due to long-term BMAL1 ablation in cardiomyocytes [3].

Compared to the drug effect in BMAL1-WT mice, the light-phase prednisone effects on myocardial tissue OCR, mitochondrial RCR, NAD^+^ and ATP levels, and *Nampt* and *Ppargc1a* expression were blunted or blocked in BMAL1-KO mice **(Fig. 3D-F)**.

Furthermore, to validate these findings in a system of complete BMAL1 knockout, we repeated this experiment and found analogous results in hearts from germline, body-wide BMAL1-WT *(Arntl^wt/wt^)* versus BMAL1-KO *(Arntl^null/null^)* littermate mice [39] at 24-hours after a pulse of light-phase prednisone versus vehicle at ZT0 **(Suppl. Fig. 3C)**. In this case, we analyzed hearts at 8 weeks of age before onset of prominent dysfunction and wasting in this strain [40].

Thus, the light-phase prednisone effects on transcriptional and metabolic remodeling of heart were cardiomyocyte-autonomous and dependent upon the drug receptor and the activating clock complex factor BMAL1.

### Chrono-pharmacology of myocardial infarction

To validate the point of circadian-dependent effects of intermittent prednisone in the context of treatment for post-MI heart failure, we compared light-phase to dark-phase prednisone regimens in MI-operated mice. Mice were randomized in treatment cohorts at 2 weeks post-MI and were treated for 6 weeks, consistent with the conditions we used earlier.

Dark-phase dosing blocked the treatment effects observed with light-phase dosing on systolic function, as shown by fractional shortening and stroke volume quantitation **(Fig. 4A)**. This correlated with analogous trends in gain of mitochondrial abundance and decreased ROS production, as assessed through mtDNA/nDNA, Mitotracker and MitoSOX assays **(Fig. 4B)**. Mass-spec profiling of left ventricle myocardium metabolites confirmed the gains of NAD^+^ and ATP content with light-phase but not dark-phase regimens. Moreover, we found that light-phase – but not darkphase – treatment increased phosphocreatine and phosphocreatine/ATP ratio, a bioenergetic parameter that is strongly affected by MI [41] and that inversely correlates with cardiovascular mortality in heart failure [42] **(Fig. 4C)**. We then assessed whether the metabolic trends correlated with pathology alleviation through molecular, histology and fibrosis markers. Unlike dark-phase treatment, light-phase treatment reduced expression of cardiac stress markers *Nppa, Nppb* and *Myh7,* as assessed through qPCR **(Fig. 4D)**. Cardiomyocyte cross-sectional area (CSA) of cardiomyocytes in the spared left ventricle was quantitated through laminin staining of tissue cryosections and showed a modest but sizeable decrease with light-phase treatment, in concordance with analogous trends with heart weight/body weight ratio. These effects were blocked with dark-phase dosing **(Fig. 4E)**. Moreover, unlike dark-phase treatment, light-phase treatment alleviated the extent of post-MI fibrotic progression, as assessed qualitatively through Masson’s staining of the border zone and quantitatively through hydroxyproline dosing in the free wall of the left ventricle **(Fig. 4F)**.

**Figure 4.**
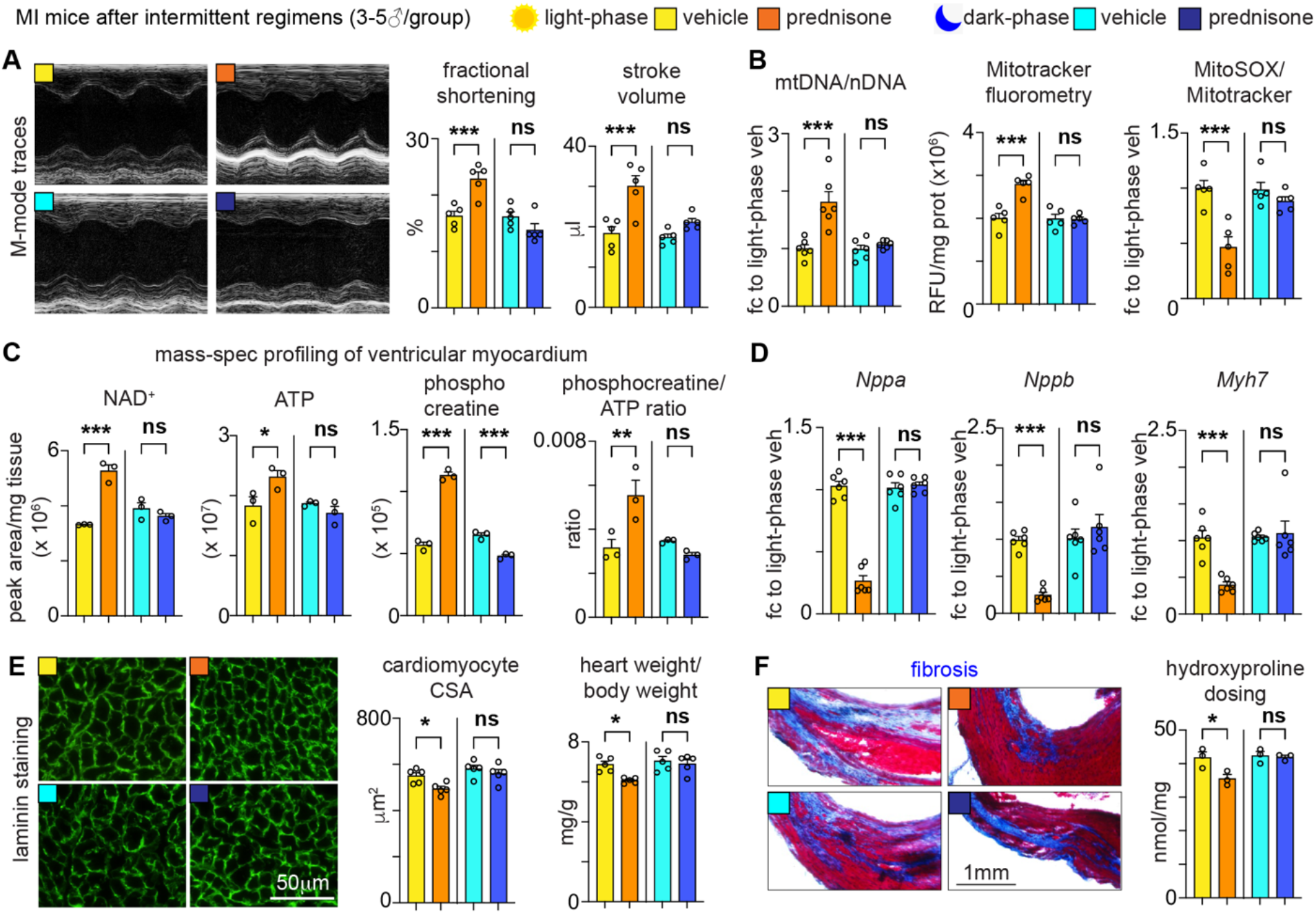
Circadian time of injection gates the intermittent prednisone effects on metabolism, function and pathology of infarcted hearts. WT mice were treated with intermittent 1mg/kg prednisone for 6 weeks from 2-weeks post-MI, with dosing at either ZT0 (light-phase treatment) or ZT12 (dark-phase treatment). **(A)** Light-phase but not dark-phase treatment improved systolic function at treatment endpoint, **(B)** correlating with time-of-dosing-specific effects on mitochondrial abundance and ROS decrease in the left ventricle myocardium. **(C)** Light-phase prednisone improved bioenergetics in the left ventricle myocardium, as indicated by improved NAD^+^, ATP and phosphocreatine/ATP levels. **(D-F)** Light-phase prednisone improved the left ventricle pathological remodeling, as indicated by decreased levels of *Nppa, Nppb* and *Myh7* expression, cardiomyocyte CSA and hydroxyproline (scar collagen) dosing. * = p < 0.05, ** = p < 0.01, *** = p < 0.001; 1w ANOVA + Sidak.

Together with our data in normal hearts, these data show that the metabolic effects of chronopharmacology with intermittent prednisone elicited functional and pathological improvements in heart failure dependent on time of dosing.

## DISCUSSION

Despite their wide usage, the cardiomyocyte-autonomous effects of glucocorticoids on heart metabolism and function remain largely unknown. Moreover, endogenous glucocorticoids are intrinsically linked to the circadian rhythm, yet little is known about time-of-intake effects on exogenous glucocorticoid action. Here we report that the exogenous glucocorticoid prednisone regulated NAD^+^, *Nampt,* ATP and *Ppargc1a* levels, as well as mitochondrial function and abundance in heart dependent on circadian time of dosing. Specifically, prednisone dosing at the start of the light-phase (ZT0) promoted those effects – i.e. during the trough of endogenous corticosterone, while dark-phase dosing (ZT12) did not. The effects were cardiac-autonomous and specific to the drug-GR axis in cardiomyocytes, as inducible ablation of cardiomyocyte-specific GR blocked the effects. Our findings are consistent with the reported cardiac dysfunction developed by mice with constitutive cardiac GR deletion [11]. In that regard, glucocorticoids are already known to improve calcium handling and contractility in cardiomyocytes [10]. The bioenergetic effects we are reporting here represent a complementary pathway of GR action in the heart. Importantly, in our inducible model, pathology was not a confounder as the acute experiment was performed immediately after ablation. Indeed, constitutive cardiac GR-knockout mice develop cardiomyopathy from 3-6 months of age [36].

In this study we used prednisone as exogenous glucocorticoid. Compared to other synthetic glucocorticoids, prednisone was effective in discriminating differences in time-of-intake likely due to short half-life and rapid uptake in striated muscles [43]. Moreover, the present data in WT, inducible GR-KO and inducible BMAL1-KO hearts are consistent with our previous findings with pro-metabolic effects of intermittent prednisone dosing in skeletal muscle dependent on circadian time-of-dosing [17]. Perhaps of importance for cardiac metabolism, here we found that the NAD^+^ increase induced by light-phase prednisone treatments correlated with decreased ROS production in normal and infarcted myocardium. Several reports have recently focused on direct effects of NAD^+^ redox state on oxidative stress in conditions of diabetic heart failure [44; 45]. Our data cannot discriminate between NAD^+^ repletion versus mitochondrial rescue as main promoter of ROS decrease downstream of light-phase prednisone, and dedicated studies will likely shed light on this pathway.

Here we did not address the role of endogenous corticosterone in mechanistically mediating or antagonizing the light-phase prednisone effects in heart. However, we did assess plasma corticosterone in our inducible cardiac GR-KO and cardiac BMAL1-KO mice and did not find differences immediately after ablation, i.e. when the control and ablated hearts presented a differential response to light-phase prednisone. However, we recognize that the role of endogenous glucocorticoids as independent variable in exogenous glucocorticoid effects on heart metabolism in vivo remains unaddressed and should be investigated through specific studies.

Our study reports on the convergence between time of prednisone administration and peripheral clock on cardiac bioenergetics. Specifically, here the convergence relied on the light-phase-restricted time of prednisone dosing. Indeed, the light-phase is the circadian phase where the activating clock complex, including BMAL1, is active in all cells including cardiac cells. Consistent with that notion of convergence, a cardiomyocyte-autonomous intact clock was required for the light-phase prednisone effects, at least acutely, as shown by the experiments with inducible cardiomyocyte-specific BMAL1 ablation and hearts from germline BMAL1-KO mice. Our data are consistent with the protective role of BMAL1 in heart, as shown by prior studies in heart function [3] and heart rate [46]. As both GR and BMAL1 are epigenetic factors regulating gene expression, dedicated studies on the effects of time-specific drug dosing on the circadian transcriptome of cardiomyocytes will be informative in dissecting transient versus adaptive changes of chronic regimens.

With regards to the overall field of “chrono-pharmacology”, we have tested here the perspective of circadian time as determinant of cardiac pharmacology outcomes of intermittent prednisone in normal and infarcted hearts. In that regard, it is important to note that circadian studies in mice are not immediately translatable to humans because of the mismatch in circadian cycles and the complexity of feeding/activity/sleep behavior in humans [47]. Nonetheless, the molecular oscillations governing heart metabolism and function are conserved [2]. Moreover, disruption of the circadian clock in the heart disrupts cardiac function in mice [3], aligning with the known cardiovascular and cardiometabolic insults induced by circadian disruption in humans [48; 49]. Thus, elucidation of molecular mediators of time-restricted pharmacology in murine models is still a cogent strategy to discern translatable mechanisms and biomarkers for the emerging clinical question of “when should you take your medicines?” [50].

Our study found notable effects on both NAD^+^ and mitochondrial respiration/abundance in myocardial tissue, suggesting a comprehensive positive effect of light-phase prednisone on aerobic energy production in heart. Moreover, the effects of light-phase prednisone on *Nampt* upregulation and NAD^+^ biogenesis in heart are consistent with the positive effects of NAD^+^ replenishment in heart failure [19; 44; 45; 51]. However, our study was performed in male mice and additional studies are required to address whether effect size and mechanisms are replicable in female mice. Furthermore, the bioenergetic effects of chronic intermittent dosing in vivo correlated with functional improvements in models of MI and diabetic cardiomyopathy. This is remarkable as our findings with intermittent (once-weekly) prednisone oppose the known correlations between once-daily regimens and heart dysfunction. Indeed, chronic once-daily glucocorticoid intake in the human population increases the chance of heart failure events in a dose-dependent manner [14]. It is therefore possible that the combination of circadian time-of-intake and dosing intermittence, at least in our experimental conditions, increased the bioenergetic effects in heart without inducing or even counteracting metabolic stress, a notion strongly supported by our prior studies in WT normal and obese mice [15; 16]. Human-relevant promising indications were recently reported by an exploratory open-label clinical trial in non-Duchenne patients with intermittent prednisone dosing at 7-9PM, which roughly corresponds to our mice injections in the early light-phase. In this 6-month-long study that compared patients at start and end of treatment, once-weekly prednisone improved performance at the 6-minute walk and 10-meters run tests without inducing the typical metabolic stress markers of chronic glucocorticoid dosing like hyperglycemia or fat mass gain [52]. Importantly for future translation, the extent to which circadian time-of-intake determines steroid effects on the immune system is still unknown and will need to be investigated. In liver, glucocorticoids induce circadian-independent effects on the immune system [60], but this still needs to be investigated in the context of heart remodeling.

In summary, this work reports cardiomyocyte-autonomous metabolic mechanisms elicited by time-specific administration of exogenous intermittent glucocorticoids. These findings pave the way to extending the chrono-pharmacology dimension to other drugs regulating metabolic function in heart and other tissues. The long-term potential of these studies is to rebalance pro-energetic versus off-target effects of long-term glucocorticoid steroid treatments and to re-evaluate glucocorticoids in the context of heart failure therapies.

## MATERIALS and METHODS

### Mice

Mice were housed in a pathogen-free facility in accordance with the American Veterinary Medical Association (AVMA) and under protocols fully approved by the Institutional Animal Care and Use Committee (IACUC) at Cincinnati Children’s Hospital Medical Center (#2020-0008). Consistent with the ethical approvals, all efforts were made to minimize suffering. Euthanasia was performed through carbon dioxide inhalation followed by cervical dislocation and heart removal.

Mice were maintained on a 12h/12h light/dark cycle. Mice were interbred on the C57BL/6 background from the following lines received from Jackson Laboratories (Bar Harbor, ME): WT C57BL/6 mice #000664; germline body-wide BMAL1-KO mice #009100; Myh6-CreERT mice #005657; GR-flox mice #021021; BMAL1-flox mice #007668.

Gene ablation was induced with tamoxifen right before start of drug treatments using a combination of i.p. (20mg/kg/day for 5 days; Sigma #T5648) and chow-mediated (40 mg/kg until 48 hours prior to start; Harlan #TD.130860) administration [53]. Analyses were performed after 2 days of tamoxifen washout.

### Prednisone administration in vivo

Single dose prednisone treatment consisted of i.p. injection of 1 mg/kg prednisone (#P6254; Sigma-Aldrich; St. Louis, MO) [54] and weekly prednisone treatment consisted of once-weekly i.p. injection of 1mg/kg prednisone. The injectable solution was diluted from a 5mg/ml stock in DMSO (#D2650; Sigma-Aldrich; St. Louis, MO) in 50μl volume. Injections were conducted either at the beginning of the light-phase (ZT0; lights-on) or at the beginning of the dark-phase (ZT12; lights-off). Tissues were harvested 24 hours after single pulse or last injection in chronic treatment. A vehicle injection of PBS (Sigma #D5652-50L) and DMSO (Fisher #BP231-100) was administered to another mouse in parallel to each prednisone injected mouse.

### Myocardial Infarction

Mice were anesthetized with briefly inhaled 2% isoflurane (Akorn 59399-106-01), intubated through the mouth, and ventilated throughout the procedure. A left lateral thoracotomy was performed and left coronary artery was identified and ligated using 8-0 prolene suture, just below the left atrium [3]. After surgery and before being placed in a recovery cage in an incubator, mice were given a single postoperative dosage of buprenorphrine-sustained release formula (3.25mg/kg body weight) by s.c. injection at 0.1 mg/kg. A sham surgical procedure was identical, but without ligature placement.

### Echocardiography

Echocardiography was performed before the MI/Sham surgery (0 week), before treatment (2 weeks) and after treatment (8 weeks). Echocardiography was performed in mice under 2% isoflurane inhalation anesthesia and the measurements were performed using a Vevo 2100 instrument with 18–38 MHz transducer (VisualSonics) as previously described [53; 55; 56], using M- and B-mode recordings for fractional shortening and stroke volume quantitation. Cardiac output was calculated as heart rate (bpm) x stroke volume (ml). Echocardiographic measurements were performed in blind.

### Mitochondrial density and NAD+-ATP targeted assays

The mtDNA/nDNA assay was performed on genomic DNA isolated using the Gbiosciences Omniprep kit (Gbiosciences #786-136). The ratio was obtained from qPCR values (absolute expression normalized to internal standard Rn45s and DNA concentration) of ND1 (mtDNA locus; primers: CTAGCAGAAACAAACCGGGC, CCGGCTGCGTATTCTACGTT) versus HK2 (nDNA locus; primers: GCCAGCCTCTCCTGATTTTAGTGT, GGGAACACAAAAGACCTCTTCTGG) [22]. For the Mitotracker-MitoSOX assay, Mitotracker Green FM powder (Invitrogen #M7514) is resuspended in 373μL of DMSO (Fisher #BP231-100) to obtain a 200μM concentration; Mitosox Red FM powder (Invitrogen #M36008) is resuspended in 13μL of DMSO (Fisher #BP231-100) to obtain a 5mM concentration. One microliter of each resuspension is added to 1mL of Mammalian Ringer’s Solution (Electron Microscopy Sciences #11763-10) containing isolated tissue from the ventricular myocardium. The solution containing myocardial tissue (~20mg cryopowder) and Mitotracker-MitoSOX is then pipetted into a 96 well plate (Corning #9017) in increments of 200μL. This plate is then read at the plate reader for fluorescence with excitation set to 490nm and emission set to 516nm for Mitotracker fluorometry, and fluorescence with excitation set to 510nm and emission set to 580nm for MitoSOX fluorescence. Mitotracker values were then normalized to protein content assayed in each well after the assay through homogenization and Bradford assay, while MitoSOX reads were normalized to Mitotracker values. NAD+-ATP targeted assays were performed on ~20mg of cryopulverized myocardial tissue (per assay) using dedicated assays: a colorimetric assay for NAD (Cayman Chemical #600480) and a luminometric assay for ATP (Cayman Chemical #700410), both then performed using a Biotek Synergy H1 Microplate Reader. All mitochondrial and metabolite dosing assays were conducted blinded to treatment groups.

### Unlabeled metabolite profiling in myocardial tissue

Total hydrophilic metabolite content was extracted from myocardial tissue at treatment endpoint through methanol: water (80:20) extraction, adapting conditions described previously [57]. Briefly, total metabolite content from myocardium was obtained from ~30mg (wet weight) tissue per animal. Frozen (−80°C) myocardial tissue was pulverized in liquid nitrogen and homogenized with ~250μl 2.3mm zirconia/silica beads (Cat # 11079125z, BioSpec, Bartlesville, OK) in 1ml metha-nol/water 80:20 (vol/vol) by means of Mini-BeadBeater-16 (Cat # 607, Biospec, Bartlesville, OK) for 1 minute. After centrifuging at 5000rpm for 5 minutes, 200μl of supernatant were transferred into a tube pre-added with 800μL of ice-cold methanol/water 80% (vol/vol). Samples were vortexed for 1 min, and then centrifuged at ~20,160 g for 15 min at 4°C. Metabolite-containing extraction solution was then dried using SpeedVac (medium power). 200μl of 50% Acetonitrile were added to the tube for reconstitution following by overtaxing for 1 min. Samples were then centrifuged for 15 min @ 20,000g, 4°C. Supernatant was collected for LC-MS analysis for Hydrophilic Metabolites Profiling as follows. Samples were analyzed by High-Performance Liquid Chromatography and High-Resolution Mass Spectrometry and Tandem Mass Spectrometry (HPLC-MS/MS). Specifically, the system consisted of a Thermo Q-Exactive in line with an electrospray source and an Ultimate3000 (Thermo) series HPLC consisting of a binary pump, degasser, and auto-sampler outfitted with a Xbridge Amide column (Waters; dimensions of 4.6 mm × 100 mm and a 3.5 μm particle size). The mobile phase A contained 95% (vol/vol) water, 5% (vol/vol) acetonitrile, 20 mM ammonium hydroxide, 20 mM ammonium acetate, pH = 9.0; B was 100% Acetonitrile. The gradient was as following: 0 min, 15% A; 2.5 min, 30% A; 7 min, 43% A; 16 min, 62% A; 16.1-18 min, 75% A; 18-25 min, 15% A with a flow rate of 400 μL/min. The capillary of the ESI source was set to 275 °C, with sheath gas at 45 arbitrary units, auxiliary gas at 5 arbitrary units and the spray voltage at 4.0 kV. In positive/negative polarity switching mode, an m/z scan range from 70 to 850 was chosen and MS1 data was collected at a resolution of 70,000. The automatic gain control (AGC) target was set at 1 × 106 and the maximum injection time was 200 ms. The top 5 precursor ions were subsequently fragmented, in a data-dependent manner, using the higher energy collisional dissociation (HCD) cell set to 30% normalized collision energy in MS2 at a resolution power of 17,500. The sample volumes of 25 μl were injected. Data acquisition and analysis were carried out by Xcalibur 4.0 software and Tracefinder 2.1 software, respectively (both from Thermo Fisher Sci-entific). Metabolite levels were analyzed as peak area normalized to total ion content and to wet tissue weight (weight before cryo-pulverization). Metabolite analysis was performed blinded to treatment groups.

### Transmission electron microscopy

Hearts of anesthetized mice were perfused with relaxing buffer (0.15% sucrose, 5% dextrose, 10 mM KCl in 1x PBS) for 3 min and then perfused with fixation buffer (1 % paraformaldehyde, 2% glutaraldehyde in 100 mM sodium cacodylate pH 7.4), then fixed overnight in fixation buffer and post-fixed in 1% OsO4 for 2h. Ultrathin sections of all tissues were counterstained with uranyl acetate and lead salts. Images were obtained using a 7600 transmission electron microscope (Hitachi) connected to a digital camera (AMT, Biosprint16). Mitochondria number and mitochondria-occupied area analyses were performed using ImageJ on 2500x images (~440μm^2^ area) blinded to treatment groups.

### Respirometry on cardiomyocytes, myocardial slices and isolated mitochondria

Basal tissue OCR values were obtained from basal rates of oxygen consumption of myocardial tissue slices at the Seahorse XF96 Extracellular Flux Analyzer platform (Agilent, Santa Clara, CA) following reported conditions [33] and as we previously detailed for tissue OCR analyses in dystrophic mice [54]. For cardiomyocytes, respirometry was performed on freshly isolated cells. Cardiomyocytes are isolated adapting previously reported conditions [58]. In anesthetized mice, the descending aorta is cut and the heart is immediately flushed by injecting 7mL pre-warmed EDTA buffer into the right ventricle. Immediately afterwards, the ascending aorta is clamped using Reynolds forceps, the heart is removed and kept in the 60mm dish with pre-warmed EDTA buffer. The digestion process is proceeded by injecting 10mL pre-warmed EDTA buffer to the apex of left ventricle. The heart is then transferred to the 60 mm dish with perfusion buffer and 3 ml perfusion buffer is injected to flush out the EDTA buffer. Then the heart is transferred to another 60 mm dish with pre-warmed 180U/ml Collagenase II buffer and 30 – 40 ml collagenase buffer is injected into the apex of LV. Then the heart is cut from the clamp, the ventricular myocardial tissue is pulled gently into small pieces using a forceps and further dissociated by gentle pipetting. The cardiomyocyte suspension is passed through a 200μm mesh, and the digestion is stopped by adding 5 ml of Stop buffer. Cardiomyocytes are then centrifuged at 100g for 3 minutes, resuspended in the stop buffer with increasing concentration of calcium (100μM, 400μM, and 1000μM), re-centrifuged and then plated in laminin-coated Seahorse plates in culture media. After 1 hour incubation the culture media is aspirated and Seahorse media is added for respirometry, which starts after an additional 1 hour equilibration in a CO_2_-free incubator. Buffer compositions: EDTA buffer contains 130mM NaCl, 5mM KCl, 0.5mM NaH_2_PO_4_, 10mM HEPES, 10mM glucose, 10mM BDM, 10mM taurine, 5mM EDTA, ph 7.8; Perfusion buffer contains 130mM NaCl, 5mM KCl, 0.5mM NaH_2_PO_4_, 10mM HEPES, 10mM glucose, 10mM BDM, 10mM taurine, 1mM MgCl_2_, ph 7.8; Collagenase buffer contains 220U/ml Collagenase II; Stop buffer consists of Perfusion buffer supplemented with 5% sterile fetal bovine serum; Culture medium (250ml) contains 2.45g Hanks’ salt, 5ml non-essential amino acids, 2.5ml MEM Vitamin Solution, 0.0875g NaHCO3, 2.5ml Pen-Strep 10X, 1 mg/ml bovine serum albumin. Regular seahorse protocol is followed with basal reads and injection of oligomycin, FCCP, Rotenone/Antimycin following reported conditions [30]. Basal OCR was calculated as baseline value (average of 3 consecutive reads) minus value after rotenone/antimycin addition (average of 3 consecutive reads). Basal OCR values were normalized to total protein content, assayed in each well after the Seahorse run through homogenization and Bradford assay. Mitochondrial ATP production rate was calculated as OCR_ATP_* 2 (mol O in mol O_2_) * 2.75 (Seahorse P/O), where OCR_ATP_ is the difference between baseline and oligomycin OCR. Nutrients: 10mM glucose or 200μM palmitate-BSA (#G7021, #P0500; Millipore-Sigma, St Louis, MO); inhibitors: 0.5μM rotenone + 0.5μM antimycin A (Agilent).

Respiratory control ratio (RCR) values were obtained from isolated mitochondria from myocardial tissue. Tissues form left ventricle are harvested from the mouse and cut up into very fine pieces. The minced tissue is placed in a 15mL conical tube (USA Scientific #188261) and 5mL of MS-EGTA buffer with 1mg Trypsin (Sigma #T1426-50MG) is added to the tube. The tube is quickly vortexed, and the tissue is left submerged in the solution. After 2 minutes, 5mL of MS-EGTA buffer with 0.2% BSA (Goldbio #A-421-250) is added to the tube to stop the trypsin reaction. MS-EGTA buffer: Mannitol-ChemProducts #M0214-45, Sucrose-Millipore #100892, HEPES-Gibco #15630-080, EGTA-RPI #E14100-50.0. The tube is inverted several times to mix then set to rest. Once the tissue has mostly settled to the bottom of the tube, 3mL of buffer is aspirated and the remaining solution and tissue is transferred to a 10mL glass tissue homogenizer (Avantor # 89026-382). Once sufficiently homogenized the solution is transferred back into the 15mL conical tube and spun in the centrifuge at 1,000g for 5 minutes at 4 degrees Celsius. After spinning, the supernatant is transferred to a new 15mL conical tube. The supernatant in the new tube is then centrifuged at 12,000g for 10 minutes at 4 degrees Celsius to pellet the mitochondria. The supernatant is discarded from the pellet and the pellet is then resuspended in 7mL of MS-EGTA buffer and centrifuged again at 12,000g for 10 minutes at 4 degrees Celsius. After spinning, the supernatant is discarded, and the mitochondria are resuspended in 1mL of Seahorse medium (Agilent #103335-100) with supplemented 5mM pyruvate (Sigma #P2256-100G) or 200μM palmitoylcarnitine (Sigma P1645-10MG), and 5mM malate (Cayman Chemical #20765). After protein quantitation using a Bradford assay (Bio-Rad #5000001), 2.5μg mitochondria are dispensed per well in 180μl total volumes and let to equilibrate for 1 hour at 37 degrees C. 20μL of 5mM ADP (Sigma #01905), 50μM Oligomycin (Milipore #495455-10MG), 100μM Carbonyl cyanide-p-trifluoromethoxyphenylhydrazone (TCI #C3463), and 5μM Rotenone (Milipore #557368-1GM)/Antimycin A (Sigma #A674-50MG) are added to drug ports A, B, C, and D respectively to yield final concentrations of 0.5mM, 50μM, 10μM, and 0.5μM. At baseline and after each drug injection, samples are read for three consecutive times. RCR was calculated as the ratio between state III (OCR after ADP addition) and uncoupled state IV (OCR after oligomycin addition). All Seahorse measurements were conducted blinded to treatment groups.

### Western Blot

Protein analysis was performed on ~50μg total lysates from heart homogenized in PBS supplemented with 1mM CaCl2, 1mM MgCl2 (#C1016, #M8266, Sigma-Aldrich; St. Louis, MO) and protease and phosphatase inhibitors (#04693232001, #04906837001, Roche, Basel, Switzerland). Blocking and stripping solutions: StartingBlock and RestorePLUS buffers (#37543, #46430, Thermo Fisher Scientific, Waltham, MA). Primary antibodies (all diluted 1:1000 for O/N incubation at +4C, all rabbit): anti-GR #A2164, anti-BMAL1 #A4714, anti-TOMM20 #A6774, anti-NCOR1 #A7046, anti-CREBBP #A17096, anti-MFN2 #A10175, anti-PARKIN #A0968 (ABclonal, Woburn, MA). Secondary antibody (diluted 1:5000 for 1-hour incubation at room temperature): donkey antirabbit (#sc-2313, Santa Cruz Biotech, Dallas, TX). Counterstain for loading control was performed with ponceau (#P7170, Sigma-Aldrich; St. Louis, MO). Blots were developed with SuperSignal Pico (cat#34579; Thermo Scientific, Waltham, MA) using the iBrightCL1000 developer system (cat #A32749; Thermo Scientific, Waltham, MA) with automatic exposure settings. WB gels and membranes were run/transferred in parallel and/or stripped for multiple antibody-based staining for densitometry analyses. Protein density was analyzed using the Gel Analysis tool in ImageJ software [59] and expressed as fold changes to control samples. All protein analyses were conducted blinded to treatment groups.

### Histological analyses

For the Masson trichrome staining, we followed general guidelines associated with the staining assay #KTMTR2LT (StatLab; McKinney, TX). Briefly, slides with tissue sections are deparaffinized through three 5 minute xylene changes and hydrated through three 5 minute absolute alcohol changes then rinsed with tap water. The slides are then covered with Bouin’s Fluid and set in a 56 degree water bath for 1.5 hours. The slides are rinsed in running tap water until tissue is colorless. Slides are placed in Weigert’s Hematoxylin (equal parts A and B) for 5 minutes. Slides are rinsed thoroughly in running tap water. Slides are placed in Biebrich Scarlet-Acid Fuchsin for 10 minutes then rinsed in distilled water. Slides are placed in Phosphomolybdic/Phosphotungstic Acid for 10 minutes. Without rinsing, slides placed in Aniline Blue Stain for 3 minutes then rinsed in distilled water. Slides are placed in 1% Acetic Acid for 3 minutes then dehydrated through 3 changes of absolute alcohol, each for 1 minute. Slides are then cleared through 3, 1 minute changes of xylene and a coverslip is placed over the sections using a permanent mounting media (Thermo Fisher 1900333).

For the cross-sectional area analysis, snap-frozen muscles are embedded in OCT and sections are cut at the cryostat then placed on slides. Using a pap pen, a hydrophobic barrier is formed around the sections. The slides are placed in a humidified container and the area of the slide containing the tissue sections is covered with permeabilization buffer (1% Triton X-100 in PBS) and left to sit at 37 degrees for 30 minutes then at room temperature for 10 minutes. The permeabilization buffer is then removed and the slides are washed with PBS. Blocking solution is then placed over the tissue sections and left to sit at room temperature for 1 hour in the same humidified container. The slides are rinsed with PBS and the primary antibody, anti-laminin (sigma L9393), is diluted 1:500 in blocking solution then placed over the tissue sections and left to sit overnight at 4 degrees. The next day, slides are washed with PBS and the secondary antibody, Alexa fluorophore 488 (Invitrogen A32790), is diluted 1:1000 in blocking solution and left to sit at room temperature for 1 hour, protected from light. Slides are rinsed with PBS and counter-stained with Hoescht, diluted 1:1000 in blocking solution, for 1 hour at room temperature, protected from light. Slides are them washed with PBS and a coverslip is placed over the slides using Prolong Gold’s Anti-fade mounting media (Cell Signaling technologies 9071S). Finished slides are stored at 4 degrees and imaged at 10X within 2 weeks. For CSA quantitation, ImageJ was used blinded to treatment groups.

### RNA-seq and qPCR analyses

RNA-seq was conducted on RNA extracted from left ventricle myocardial tissue. Total RNA was extracted as detailed above and re-purified using the RNeasy Mini Kit (Cat #74104; Qiagen, Germantown, MD). RNA-seq was performed at the DNA Core (CCHMC) blinded to treatment groups. 150 to 300 ng of total RNA determined by Qubit (Cat#Q33238; Invitrogen, Waltham, MA) high-sensitivity spectrofluorometric measurement was poly-A selected and reverse transcribed using Illumina’s TruSeq stranded mRNA library preparation kit (Cat# 20020595; Illumina, San Diego, CA). Each sample was fitted with one of 96 adapters containing a different 8 base molecular barcode for high level multiplexing. After 15 cycles of PCR amplification, completed libraries were sequenced on an Illumina NovaSeqTM 6000, generating 20 million or more high quality 100 base long paired end reads per sample. A quality control check on the fastq files was performed using FastQC. Upon passing basic quality metrics, the reads were trimmed to remove adapters and low-quality reads using default parameters in Trimmomatic [60] [Version 0.33]. The trimmed reads were then mapped to mm10 reference genome using default parameters with strandness (R for single-end and RF for paired-end) option in Hisat2 [61] [Version 2.0.5]. In the next step, transcript/gene abundance was determined using kallisto [62] [Version 0.43.1]. We first created a transcriptome index in kallisto using Ensembl cDNA sequences for the reference genome. This index was then used to quantify transcript abundance in raw counts and counts per million (CPM). Differential expression (DE genes, FDR<0.05) was quantitated through DESeq2 [63]. PCA was conducted using ClustVis [64]. Gene ontology pathway enrichment was conducted using the Gene Onthology analysis tool [65].

For RT-qPCR assays, total RNA was extracted from cryo-pulverized myocardial tissue with Trizol (#15596026; Thermo Fisher Scientific, Waltham, MA) and 1mg RNA was reverse-transcribed using 1X qScript Supermix (#95048; QuantaBio, Beverly, MA). RT-qPCRs were conducted in three replicates using 1X Sybr Green Fast qPCR mix (#RK21200, ABclonal, Woburn, MA) and 100nM primers at a CFX96 qPCR machine (Bio-Rad, Hercules, CA; thermal profile: 95C, 15sec; 60C, 30sec; 40X; melting curve). The 2-ΔΔCT method was used to calculate relative gene expression. was used to calculate the relative gene expression (Schmittegen and Livak, 2008). GAPDH was used as the internal control. Primers were selected among validated primer sets from the MGH PrimerBank; IDs: Fkbp5 (56753884a1), Tsc22d3 (11907580a1), Nampt (10946948a1), Ppargc1a (6679433a1), Ppara (31543500a1), Rora (1869971a1), Clock (21950741a1), Arntl (4586560a1), Gapdh (6679937a1).

### Statistics

Statistical analyses were performed using Prism software v8.4.1 (Graphpad, La Jolla, CA). The Pearson-D’Agostino normality test was used to assess data distribution normality. When comparing two groups, two-tailed Student’s t-test with Welch’s correction (unequal variances) was used. When comparing three groups of data for one variable, one-way ANOVA with Sidak multi-comparison was used. When comparing data groups for more than one related variable, two-way ANOVA was used. For ANOVA and t-test analyses, a P value less than 0.05 was considered significant. When the number of data points was less than 10, data were presented as single values (dot plots, histograms). Tukey distribution bars were used to emphasize data range distribution in analyses pooling larger data point sets per group (typically > 10 data points). Analyses pooling data points over time were presented as line plots connecting medians of box plots showing distribution of all data per time point.

## ACKNOWLEDGMENTS

RNA-seq analyses were conducted at the Cincinnati Children’s DNA Core. Metabolomic analyses were conducted at the Northwestern University Metabolomics Core. We are grateful to Dr. J. Hogenesch (CCHMC), Dr. K. Yutzey (CCHMC), Dr. K. Esser (UF) and Dr. K. Lamia (Scripps) for helpful discussions. This work was supported by the following grant support: NIH grant DK121875 (MQ), NIH grant HL158531 (MQ), CCHMC Trustee Award (MQ), CCHMC Heart Institute Translational Grant (MQ).

---

**Supplementary Figure 1.**
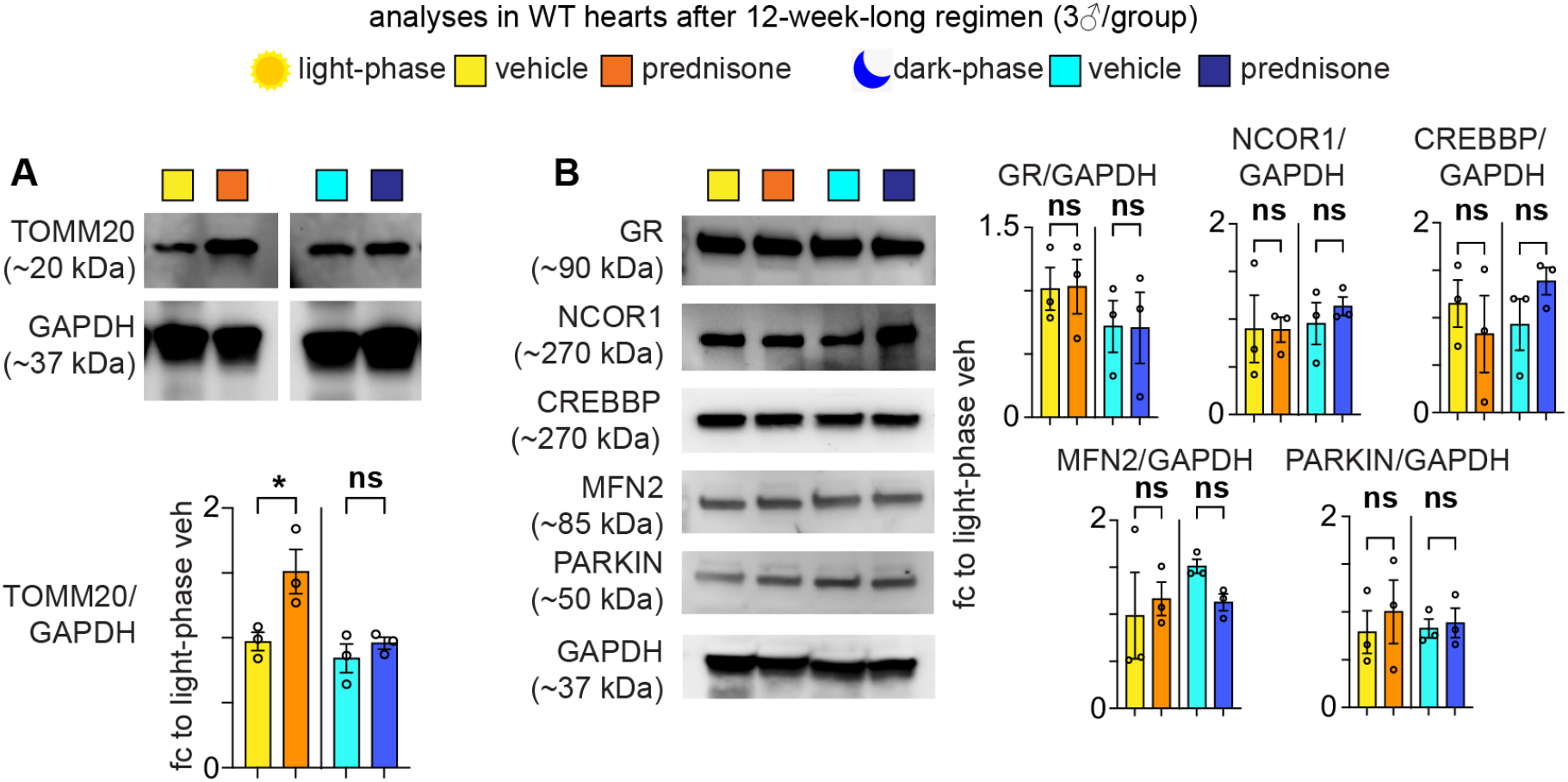
Additional data related to Figure 1. **(A)** Time-specific effects of intermittent prednisone on myocardial mitochondrial abundance were indicated also by TOMM20 protein levels. **(B)** No regimen-specific changes in protein levels of GR, its co-factors NCOR1 and CREBBP, or mitochondrial remodeling factors MFN2 and PARKIN were found. * = p < 0.05, ** = p < 0.01, *** = p < 0.001; 1w ANOVA + Sidak.

**Supplementary Figure 2.**
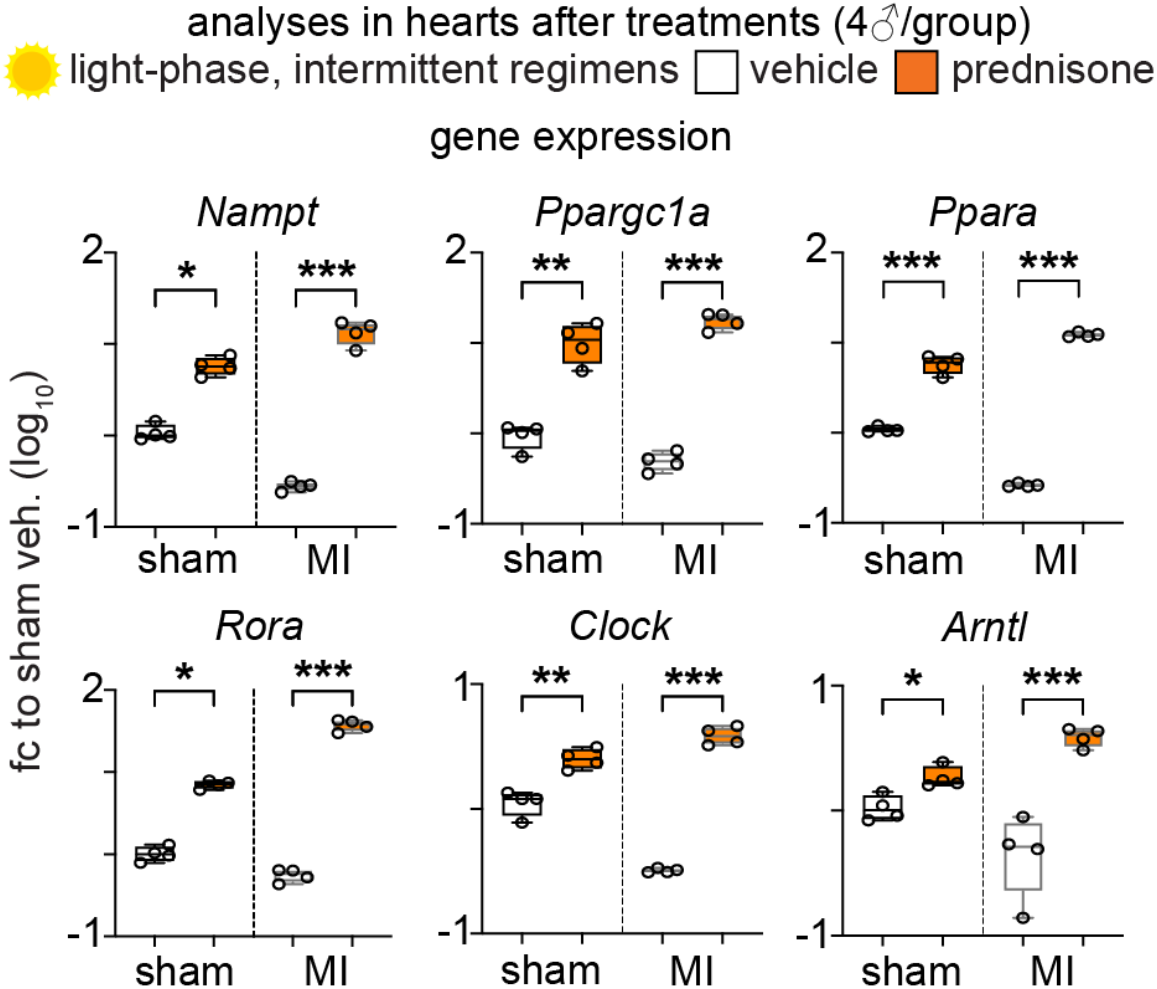
Additional data related to Figure 2. Validation of concordant genes highlighted by RNA-seq profiling after light-phase intermittent prednisone. qPCR analyses confirmed treatment-driven upregulation of clock and metabolic genes in both sham and MI hearts as compared to respective vehicle controls. * = p < 0.05, ** = p < 0.01, *** = p < 0.001; 1w ANOVA + Sidak.

**Supplementary Figure 3.**
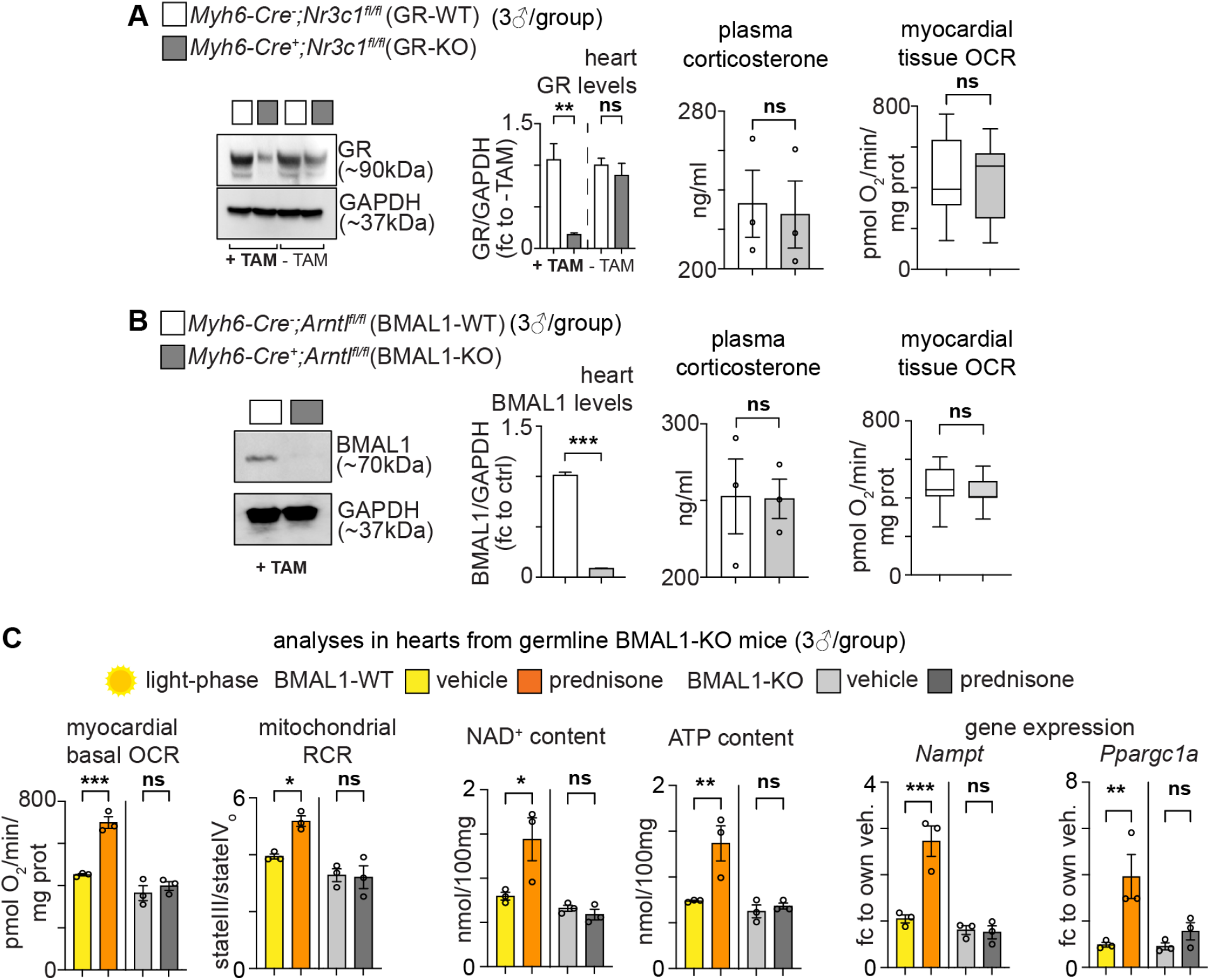
Additional data related to Figure 3. **(A)** GR was ablated for ~85% of its WT levels after tamoxifen treatment and washout at 8 weeks of age. Immediately after cardiac-restricted ablation and washout, GR-WT and GR-KO mice did not show significant changes in plasma corticosterone or myocardial mitochondrial respiration at baseline. **(B)** Similar ablation and baseline trends were found after cardiomyocyte-specific inducible BMAL1 ablation. **(C)** Analogous results to the ones obtained with inducible ablation of BMAL1 were found in hearts from germline BMAL1-KO mice. * = p < 0.05, ** = p < 0.01, *** = p < 0.001; 1w ANOVA + Sidak.

**Supplementary Table 1** – List of genes with >2 fold-change variation and >10CPM abundance found with treatment versus vehicle comparisons in sham and MI hearts.

**Supplementary Table 2** – List of GO terms enriched from DE genes in sham and MI heart comparisons.

## Notes

**Conflicts of interest** – MQ is listed as co-inventor on a patent application related to intermittent glucocorticoid use filed by Northwestern University (PCT/US2019/068618). All other authors declare they have no competing interests.

### Competing Interest Statement

MQ is listed as co-inventor on a patent application related to intermittent glucocorticoid use filed by Northwestern University (PCT/US2019/068618). All other authors declare they have no com-peting interests.

### Summary of Updates

New cardiomyocyte-specific transgenic models and new in vivo chrono-pharmacology analyses have been added.

